# The role of the ventral midline thalamus in the retrieval of precise temporal memories

**DOI:** 10.64898/2026.05.11.724442

**Authors:** Alcides P. Lorenzo Gonzalez, Timothy Allen

## Abstract

Interval timing (IT) is the ability to time events in the range from seconds to a few minutes, allowing animals to organize behavior in time at short durations. IT relies on two cognitive functions: 1) Measuring the passage of time; 2) Storing and retrieving temporal memories in a context appropriate manner. The hippocampus (HC) and medial prefrontal cortex (mPFC) have been shown critical to the accuracy and precision of time-contingent instrumental responses in IT. The anatomy supporting mPFC-HC interactions, required for memory encoding and retrieval, include projections from HC to mPFC, and indirect bidirectional connections through the ventral midline thalamus (VMT), most notably reuniens. Here, we explored VMT’s role in retrieving fixed-interval (FI) temporal memories. Rats were trained on a 5s FI signaled by an auditory cue and demonstrated temporal memory by poking predominantly at the time of the expected reward. Timing responses on individual trials were classified into on-time, early, and random response. Across sessions, random response trials decreased following training. Next, we switched training to longer intervals (20s or 80s; daily sessions for weeks). To probe the role of the VMT in temporal memory retrieval, we infused the GABA_A_-agonist muscimol, or saline, before training sessions. Results show that VMT muscimol infusions decreased timing precision. Also, at both intervals, the number of on-time response trials decreased, and the number of random response trials significantly increased. The number of early response trials had no significant change at 20s, and significantly decreased at 80s. Overall, our results suggest that the VMT is critical for precise retrieval of temporal memories. We also describe per-trial response patterns with characteristics consistent across all trained intervals, suggesting multiple behavioral strategies at play during interval timing.

## INTRODUCTION

The accurate timing of events in the range of seconds to a few minutes, which we refer to as interval timing, allows animals to organize behavior in time at short durations (Brunner et al., 1992; Pyke et al., 1977). In mental health disorders such as schizophrenia, attention-deficit disorder, among others, patients display symptoms related to a diminished timing ability (Bonnot et al., 2011; Lee et al., 2009; Penney et al., 2005; Rubia et al., 2009). In the brain, the timing of intervals makes use of two fundamental functions. First, the ability to measure the passage of time: a striatum-dependent function that relies on an “internal clock” and is sensitive to somatosensory and proprioceptive cues (Buhusi & Meck, 2005; Drew et al., 2007; Gouvea et al., 2015; Meck, 2006; Mello et al., 2015; Paton & Buonomano, 2018); and second, the ability to correctly retrieve these time memories in a context appropriate manner (De Corte et al., 2022).

Two brain areas widely associated with memory, hippocampus (HC) and medial prefrontal cortex (mPFC), have been shown to play a role in interval timing. Damage or inactivation of HC reduces accuracy of interval timing by producing and underestimation of the target interval (Meck, 1988; Meck et al., 1984; Tam & Bonardi, 2012a, 2012b; Tam et al., 2013; Yin & Meck, 2014). Damage or inactivation of mPFC reduces precision by reducing the number of timing-contingent instrumental responses (Buhusi et al., 2018; Dietrich & Allen, 1998; Dietrich et al., 1997; Elcoro et al., 2014; Kim et al., 2009). Interactions between HC and mPFC are required for memory functions (Floresco et al., 1997; Benchenane et al., 2010), specifically during memory encoding and retrieval (Churchwell et al., 2010). These interactions are supported by anatomical connections as mPFC receives inputs from HC (Hoover & Vertes, 2007; Jay & Witter, 1991), however, there is no direct return from mPFC into HC (Vertes, 2004), which suggests an indirect route of communication. The ventral midline thalamus (VMT) sits at the anatomical nexus between the HC and mPFC by projecting bi-directionally to both areas (Hoover & Vertes, 2007; Vertes, 2002; Vertes et al., 2006; 2007; Varela et al, 2014; Viena et al. 2022). This thalamic structure can support the communication between mPFC and HC; thus, it can play a role in the memory functions associated to timing.

The VMT includes the reuniens (RE) and rhomboid (Rh) nuclei. Studies suggest that RE projections to mPFC and HC participate in spatial navigation (Ito et al., 2015, Viena et al, 2018), navigational strategy shifting (Cholvin et al., 2013), consolidation of spatial (Loureiro et al., 2012) and sequence memories (Jayachandran et al., 2019), acquisition of associative learning (Eleore et al., 2011), spatial working memory (Hembrook et al., 2012), as well as the processing of contextual information being indirectly transmitted from mPFC to HC (Xu & Sudhof, 2013). RE seems to be particularly important during working memory tasks (Hallock et al., 2013), more specifically when memory retrieval is delayed in time (Hallock et al., 2016; Viena et al., 2018).

In addition to its anatomical and behavioral relationship, research shows that neural activity in RE/Rh participates in the processing of information originating in mPFC and HC. VMT projections are a prominent direct source of excitatory inputs into mPFC and HC (Di Prisco & Vertes, 2006) where they can produce neural facilitation. Low frequency oscillatory activity in RE can modulate oscillation amplitude in HC possibly by the activation of interneurons (Dolleman-van der Weel et al., 1997, 2017), modulation that seems to affect spatial working memory (Duan et al., 2012). Systemic ketamine injections produce a characteristic increase in hippocampal delta activity that gets blocked by RE inactivation (Zhang et al., 2012) pointing to thalamic control over activity in HC. Conversely, neurons in RE have been shown to display heterogeneous responses to HC oscillatory activity (Lara-Vasquez et al., 2016). Ferraris et al., (2018) have shown that RE neurons increase in activity precedes gamma bursts that synchronize HC and mPFC during slow wave activity. Although synchronization at higher oscillatory frequencies in the mPFC-RE-HC circuit has been observed during sequence memory retrieval (Jayachandran et al., 2023), a systematic characterization of low-frequency oscillatory activity dynamics in RE/Rh during memory tasks is still required. Understanding the role of the VMT in interval timing and context-appropriate temporal memory retrieval can provide insight into how the brain handles these cognitive functions, as well as shed light into the role of thalamic activity controlling mPFC-HC communication during learning and memory.

## METHODS

### Subjects

Eight male Long-Evans rats (Charles River Laboratories) weighing approximately 300 g upon arrival were used in these experiments. Rats were individually housed and maintained on a 12 h inverse light/dark cycle. Rats training on the interval timing task had ad libitum access to food and enrichment, water consumption was restricted to 5 minutes daily. Water restriction started one week before beginning behavioral training, it was discontinued 5 days before implant surgery, and reinstated one week post-op to ensure proper recovery. All training sessions were conducted during the dark phase (active period) of the light cycle. All experimental procedures using animals were conducted in accordance with the Florida International University Institutional Animal Care and Use Committee (FIU IACUC).

### Interval timing task

The behavioral task used consists of a modified Fixed-Interval task in which rats are trained in each trial to nose-poke for a sugar water reward contingent on the length of a simultaneous auditory stimulus (white noise). The start of each trial is marked by the beginning of the auditory cue, and the end occurs upon the first nose-poke after the predetermined time interval has elapsed. In the case that no nose-pocks trigger the reward, the auditory cue remains on for another 40 seconds, and the trial is marked as failed. Intertrial intervals were assigned a random duration between 30-50 s. Rats demonstrated interval timing memory by poking predominantly at the time of the expected reward.

### Interval timing apparatus

The behavioral apparatus consisted of an enclosed end of a linear track that allowed the rat to easily turn around and explore the area. The nose port consisted of a hole in the maze that contained a photobeam sensor used to detect nose port entries. The water port was located right below the nose port and was used for reward delivery. A wired speaker was located two feet above and facing the maze and served as the source of the auditory cue. An OmniPlex Neural Recording Data Acquisition System (Plexon, TX) and digital input/output devices (National Instruments, TX) were used to detect all event times and electrically control the hardware. All aspects of the task were automated using custom scripts written in MATLAB (MathWorks R2016a). Cineplex behavior software (Plexon, TX) was used to video record all the behavioral sessions.

### VMT cannula implant

The cannula implants were designed and 3D printed in the laboratory in accordance with the RatHat patented model (Allen et al., 2020). A 26G stainless steel hypodermic tube (A-M Systems, WA) was cemented to the printed model at a 15° angle and used as guide cannula. The infusion cannula was made using a 32G stainless steel hypodermic tube (A-M Systems, WA).

### Implant surgery

Rats were anesthetized with isoflurane (5% induction, 2-3% maintenance) mixed with oxygen (800 mL/min) and placed in a stereotaxic apparatus (David Kopf Instruments, Model 900). The scalp was shaved and sterilized using 70% isopropyl alcohol and Betadine. Ophthalmic ointment was applied to the eyes for protection, and a thermal probe was inserted rectally for temperature monitoring. A burr hole was drilled over the midline thalamus and the custom-built cannula implant was inserted and placed above the reuniens and rhomboid nuclei of the ventral midline thalamus (−2.0/-2.1 mm AP; +2.0 mm L; 6.5 – 7.0 mm DV). Bone screws were anchored to the skull, and the screws and cannula implant were affixed to the skull using dental cement. Dummy cannulas (7 mm in length) were inserted to cover the implanted guide and prevent any insults.

### Muscimol infusions

Rats were carefully immobilized using a swaddle. The custom-built infusion cannula was inserted 8 mm into the guide cannula, and 0.5 ul of muscimol were infused at a rate of 9 nL/s using a Nanoject III Automatic Nanoliter Injector (Drummond, PA). All eight rats were initially infused with a muscimol concentration of 0.125 mg/mL, one rat did not survive the first session, and two others appeared lethargic during behavior. Their muscimol concentration was halved (0.063 mg/mL) which allowed them to perform during behavior. Upon finishing the procedure, the infusion cannula was left in place for 5 min to allow for drug diffusion and to prevent backwash during removal. The infused rat was placed back in its home cage for 30 min before behavioral training. If drug or saline infusions were unsuccessful, the guide cannula was assumed clogged, thus a 7.5 mm long dummy cannula was inserted to clear the guide cannula. This was the case for two rats, which later received multiple successful infusions.

### Histology

After completing all experiments, rats were transcardially perfused with 200 ml heparinized phosphate-buffered saline, followed by 200 ml of 4% paraformaldehyde (pH 7.4; MilliporeSigma). Brains were post-fixed overnight in 4% paraformaldehyde and placed in a 30% sucrose solution for cryoprotection. Frozen brains were cut in the coronal plane on a Leica CM3050 S cryostat in 40 uM sections. Sections were mounted, Nissl-stained, and coverslipped with Permount to visualize cannula tracks and lesions. The sections were then mapped onto coordinate from the Paxinos and Watson Rat Brain Atlas (2013). Drug diffusion in tissue was estimated based on the infused volume.

### Data analysis

Plexon data was preprocessed to generate binned (1s bins) timestamps of nose-pokes, and auditory cue ON/OFF status per trial. To generate the response rate curves, all trials in a session were aligned by the start of the cue. From these curves we extracted the magnitude and time of peak response, the mean cued and pre-cue (20s preceding cue start) response rates, as well as total pokes per session. These behavioral metrics were averaged first by session, and then by subject. To account for individual differences in poking, response rates were normalized using the mean peak response per animal of the last week of training for the trained interval. For our clustering analysis we created an array containing all non-treatment trials from every rat and used a kmeans clustering algorithm to separate by whole-trial response rate. The obtained cluster centroids where then used to cluster trials from the treatment sessions. All statistical significance was tested assuming non-normal data. The non-parametric Wilcoxon Signed Rank test was used to compare paired behavioral data for drug, saline, and control groups. The non-parametric Wilcoxon Ranksum was used to compare clustered behavioral data.

## RESULTS

### 5s interval timing

To assess the ability of rats to measure intervals of time, we initially trained them to nose-poke in expectation of a sugar water reward. Next, we trained them daily using a modified 5s FI schedule with an auditory cue signaling the duration of the trial (Figure 1A). Rats were able to nose-poke at any time and were only rewarded for the first poke after the trained interval had elapsed, which would also turn off the auditory cue. The initial 5s IT session was characterized by nose-poking at random, displaying a flat response curve with a small dip at the time of reward, as the animal stopped poking to consume the reward. Across a few days responses started to accumulate within the cued interval; after two weeks of training their response were occurring mostly within the cued time, increasing in expectancy of reward, peaking at the trained 5s interval, and with minimal pre-cue responses (Figure 1B). The total number of responses per session didn’t significantly change throughout the weeks of training, suggesting that the total behavioral output of the rats was not diminished but reallocated through conditioning with the auditory cue, the timing of the interval, and the water reward.

**Figure 1.**
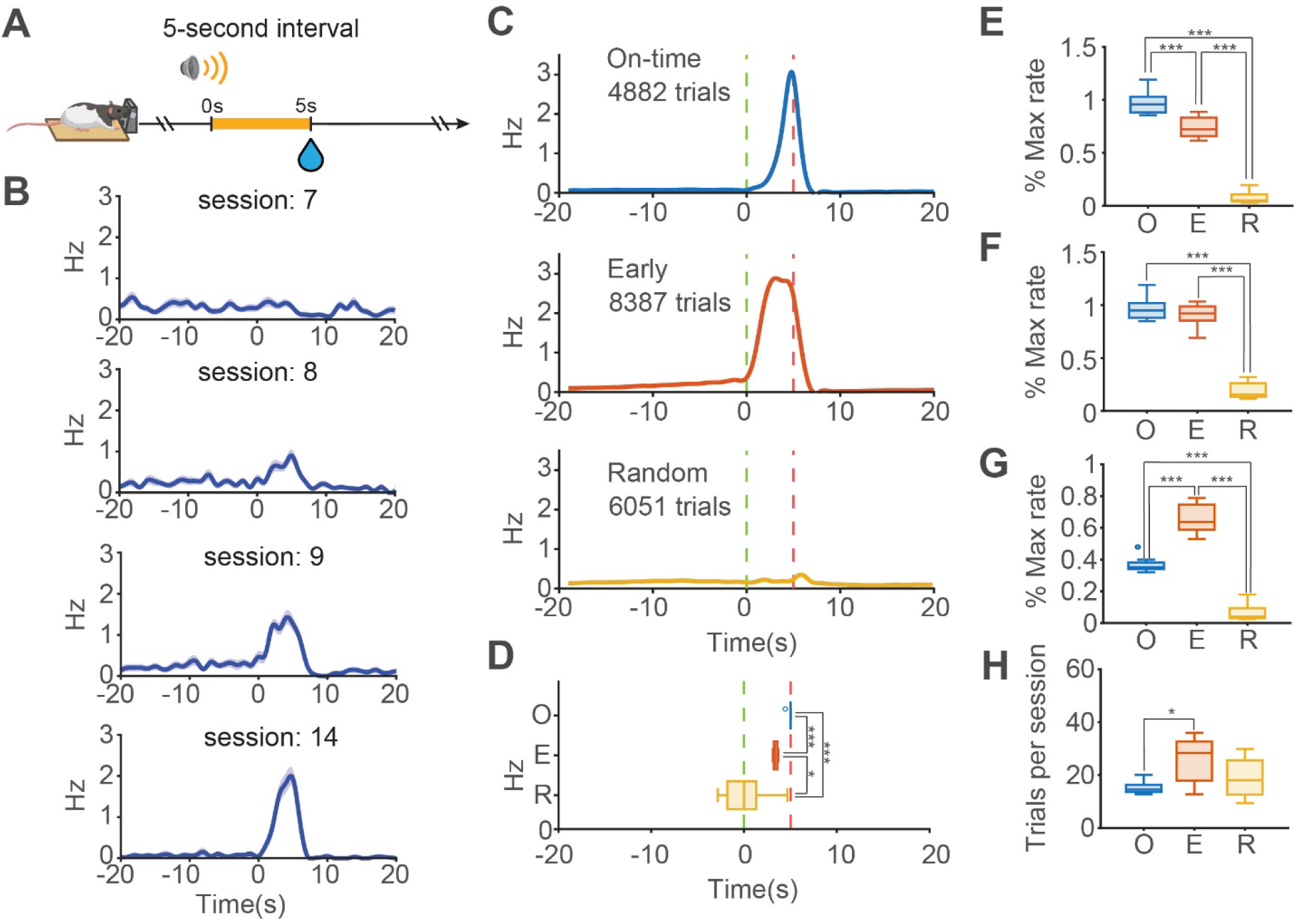
Timing at 5s interval. **A.** Schematic of the IT task using a 5s interval. **B.** Example session average response curve for 5s IT learning. Session 7 shows the first 5s IT session. Across days responses accumulate around the trained interval. **C.** Response clusters from all 5s IT trials. Cluster name is based on the precision of the timing in the response curve. **D.** On-Time Response (O) peaks at the time of expected reward; Early Timing Response (E) peaks before the expected reward; Random Timing Response (R) peaks earlier than both OTR and ETR. Each cluster is characterized by a pattern of response rate. *Cluster metrics:* **E.** Response rates at time of interval. **F.** Peak response rates. **G.** Mean cued response rates. **H.** Average number of trials per session.

The average response rate curves for 5s IT showed a specific pattern that pointed to multiple distribution peaks within the mean curve. To further understand this observation, we aggregated all trials per time interval and converted them into vectors of normalized response rate. These vectors of size 40 included the trained 5s interval, 20s pre-cue, and 15s post trained interval. We ran a k-mean clustering algorithm on these vectors, and determined the optimal number of clusters to be 3. Responses per trial were then clustered into three categories characterized by the observed times of peak response. We labeled these clusters as On-Time Response (OTR), Early Timing Response (ETR), and Random Timing Response (RTR) (Figure 1C).

OTR trials displayed an average response that peaked approximately at the time of reward (µ = 4.9s, σ = 0.19s). ETR trials showed a peak response rate before the expected time of reward (µ = 3.4s, σ =0.23s). RTR trials showed a high variability in peak response time, closer to cue onset (µ = 0.12s, σ = 2.5s) (Figure 1D). Besides time of peak response rate, other quantitative features helped describe these behavioral clusters. Response rates at the time of expected reward were significantly different between all three clusters (OTR: µ = 0.97, ETR: µ = 0.74, RTR: µ = 0.077; Wilcoxon ranksum: p < 0.001) (Figure 1E). OTR and ETR trials had an average peak response rate (normalized to % of maximal response – see Methods) significantly higher than RTR trials (OTR: µ = 0.97, ETR: µ = 0.91, RTR: µ = 0.19; Wilcoxon ranksum: p < 0.001) (Figure 1F). ETR trials showed a mean cued response rate higher than OTR (OTR: µ = 0.37, ETR: µ = 0.66; Wilcoxon ranksum: p < 0.001) as the change in response curve slope occurs earlier in the ETR trials. Conversely, RTR trials were characterized by a very low mean cued response rate (RTR: µ = 0.067) (Figure 1G). Pre-cue mean response rates were not significantly different between clusters (OTR: µ = 0.016, ETR: µ = 0.029, RTR: µ = 0.027). Lastly, each of the response clusters constituted roughly a third of all trials per session with ETR trials having a slight advantage over OTR trials (OTR: µ = 15.2 trials, ETR: µ = 25.8 trials; Wilcoxon ranksum: p = 0.0201; RTR: µ = 18.9 trials) (Figure 1H). These consistent differences across trial types suggest that the behavior displayed during interval timing is not simply a continuum of variable responses around an estimated interval. More specifically, they suggest that rats could be using multiple behavioral strategies in expectation of a timed reward, and that these strategies can be masked by session averaging of responses.

### Cannula implant

Following several weeks of 5s IT training, rats were surgically implanted with a guide cannula targeting the VMT (Figure 2). Rats were allowed to recover for one week and re-trained in 5s IT for an extra week. Post-implanting all rats still demonstrated interval timing and remained engaged with the behavioral task. All implanted rats were then moved on to training a 20s interval using the same behavioral protocol (Figure 3A).

**Figure 2.**
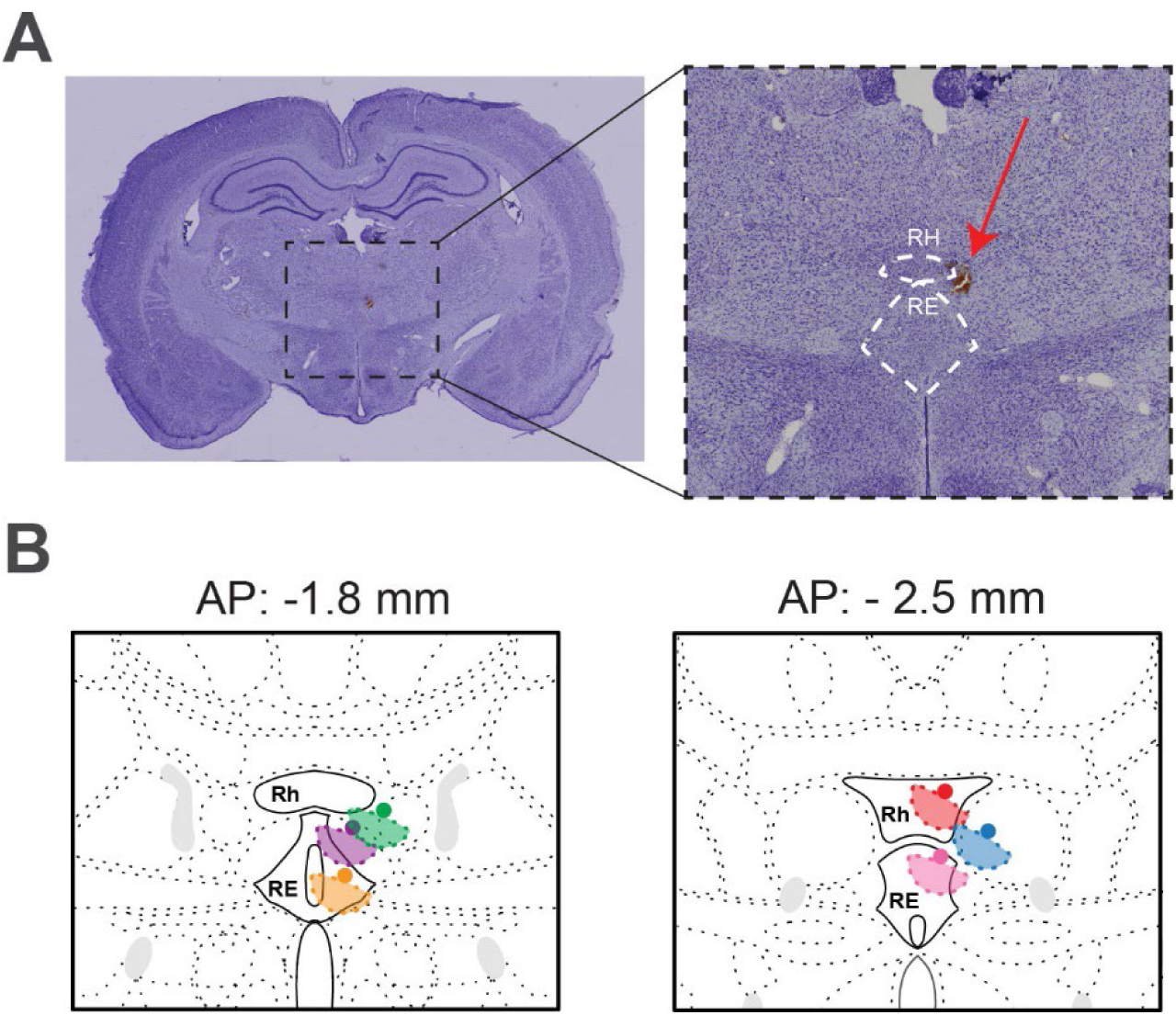
Infusion cannula placement targeting VMT. **A.** Example coronal section showing cannula placement above VMT (red arrow). **B.** Schematic of all cannula placements for all experimental rats.

**Figure 3.**
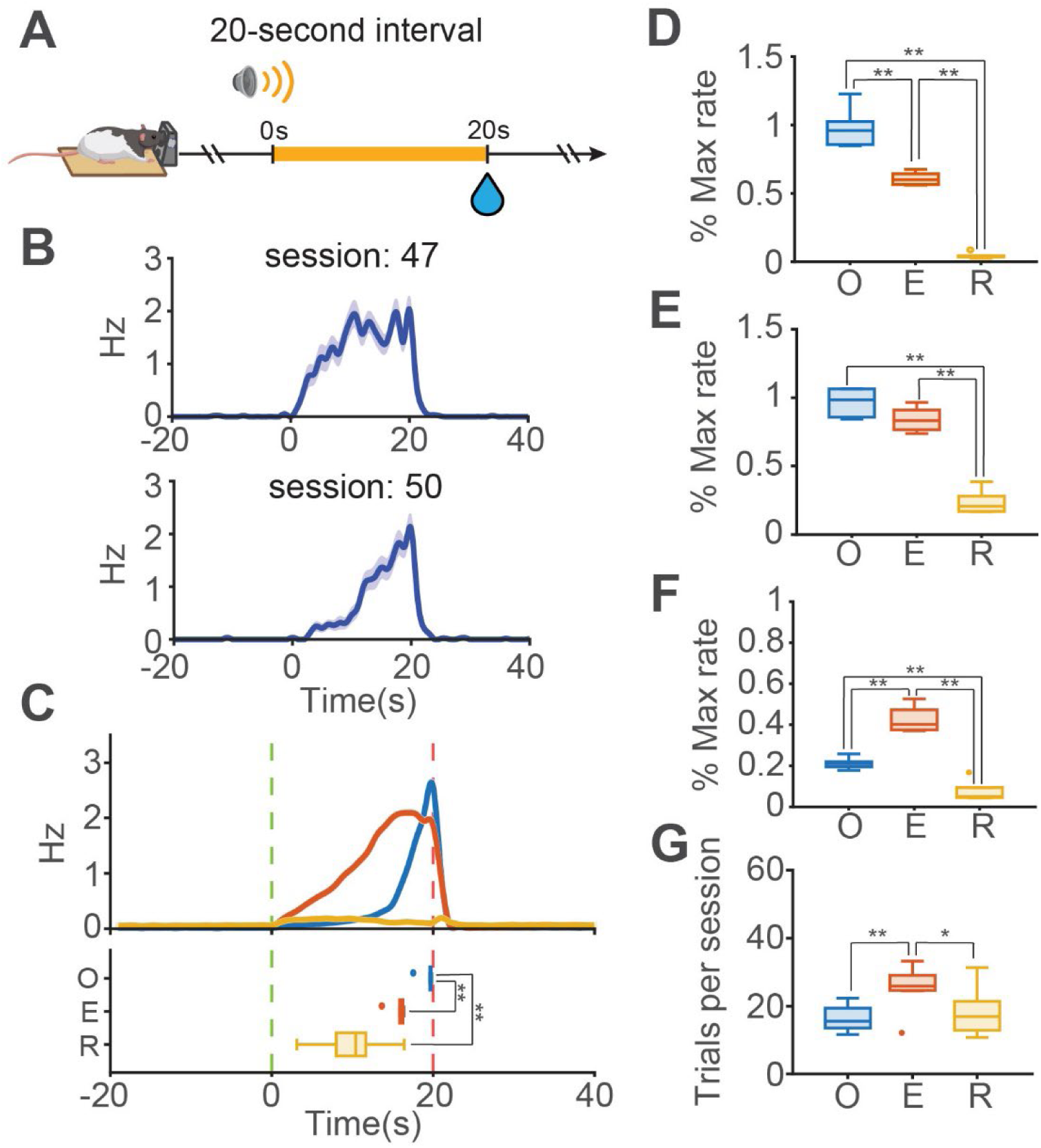
Timing at 20s interval. **A.** Schematic of the IT task using a 20s interval. **B.** Example session average response curves for 20s IT learning. Session 47 shows the first 20s IT session. **C.** (Top) Response clusters from all 20s IT trials. (Bottom) Time of peak response per cluster. On-Time Response (O) peaks at the time of expected reward; Early Timing Response (E) peaks before the expected reward; Random Timing Response (R) peaks earlier than both OTR and ETR. As in 5s IT each cluster is characterized by a pattern of response rate. *Cluster metrics:* **D.** Response rates at time of interval. **E.** Peak response rates. **F.** Mean cued response rates. **G.** Average number of trials per session.

### 20s interval timing

Due to the familiarity with the task, the first 20s session did not show the random nose-poking pattern seen at 5s IT. The curve of the new trained interval now began at cue onset with very few pre-cue responses (Figure 3B). A week later, response rates were lower during the first 5s of the auditory cue, and peaked at 20s. Within single sessions we observed the response pattern narrowing closer to the expected reward time across trials, showing how learning also occurred during a training session. These observations confirmed that rats were learning the trained time, rather than uniquely responding to the auditory cue.

To probe the presence of the response clusters beyond 5s IT, we clustered the 20s IT response trials in three clusters using a vector size of 60 this time (20s interval + 20s pre-cue + 20s post interval). All three characteristic clusters observed at 5s IT were also present at this longer interval. OTR trials peaked approximately at the time of reward (µ = 19.4s, σ = 0.89s), ETR trials peaked before the expected time of reward (µ = 15.7s, σ =1.01s), and RTR trials displayed a larger variability in peak response time (µ = 9.9s, σ = 4.4s) (Figure 3C). Response rates at the time of expected reward were significantly different between all three clusters (OTR: µ = 0.98, ETR: µ = 0.61, RTR: µ = 0.044; Wilcoxon ranksum: p = 0.0022) (Figure 3D). The average peak response rate in OTR and ETR trials was significantly higher than in RTR trials (OTR: µ = 0.97, ETR: µ = 0.84, RTR: µ = 0.24; Wilcoxon ranksum: p = 0.0022) (Figure 3E). Similar to the 5s IT clusters, 20s IT trials showed a significantly higher mean response rate in the ETR vs OTR cluster (OTR: µ = 0.21, ETR: µ = 0.42; Wilcoxon ranksum: p = 0.0022). Mean response rate for RTR trials were significantly lower than both OTR and ETR trials (RTR: µ = 0.075; Wilcoxon ranksum: p = 0.0022) (Figure 3F). Lastly, the number of trials in the ETR cluster was significantly higher than both in the OTR and RTR clusters (OTR: µ = 15.5 trials, ETR: µ = 26.7 trials; Wilcoxon ranksum: p = 0.0087; RTR: µ = 18.9 trials; Wilcoxon ranksum: p = 0.041) (Figure 3G).

At both 5s and 20s IT the quantitative differences between response trial clusters were consistent supporting the conclusion of multiple coexistent behavioral strategies at display the interval timing task, that scale up with the timed interval. This scaling property is also supported by ETR trials becoming the majority (insofar as early responses can be looked at as timing errors).

### VMT inactivation in 20s IT

After several weeks of 20s interval timing training we began a low concentration pre-session infusion of muscimol (0.125 mg/mL) targeting the VMT. The days of drug infusion were followed by a day of no infusion, a day of saline infusion, and another day of no infusion; thus, a minimum of three days post muscimol treatment allowed us to control for any lingering drug effects. The response curve during VMT inactivation sessions showed two main characteristics: increased pre-cue response rates (muscimol: µ = 0.14, no infusion: µ = 0.011, saline: µ = 0.016) and decreased peak response rates (muscimol: µ = 0.56, no infusion: µ = 1.00, saline: µ = 1.007). When compared to the saline and no infusion sessions, both rate changes were statistically significant (Wilcoxon sign-ranked: p = 0.031) (Figure 4A-D) suggesting a decrease in the precision of the temporal response.

**Figure 4.**
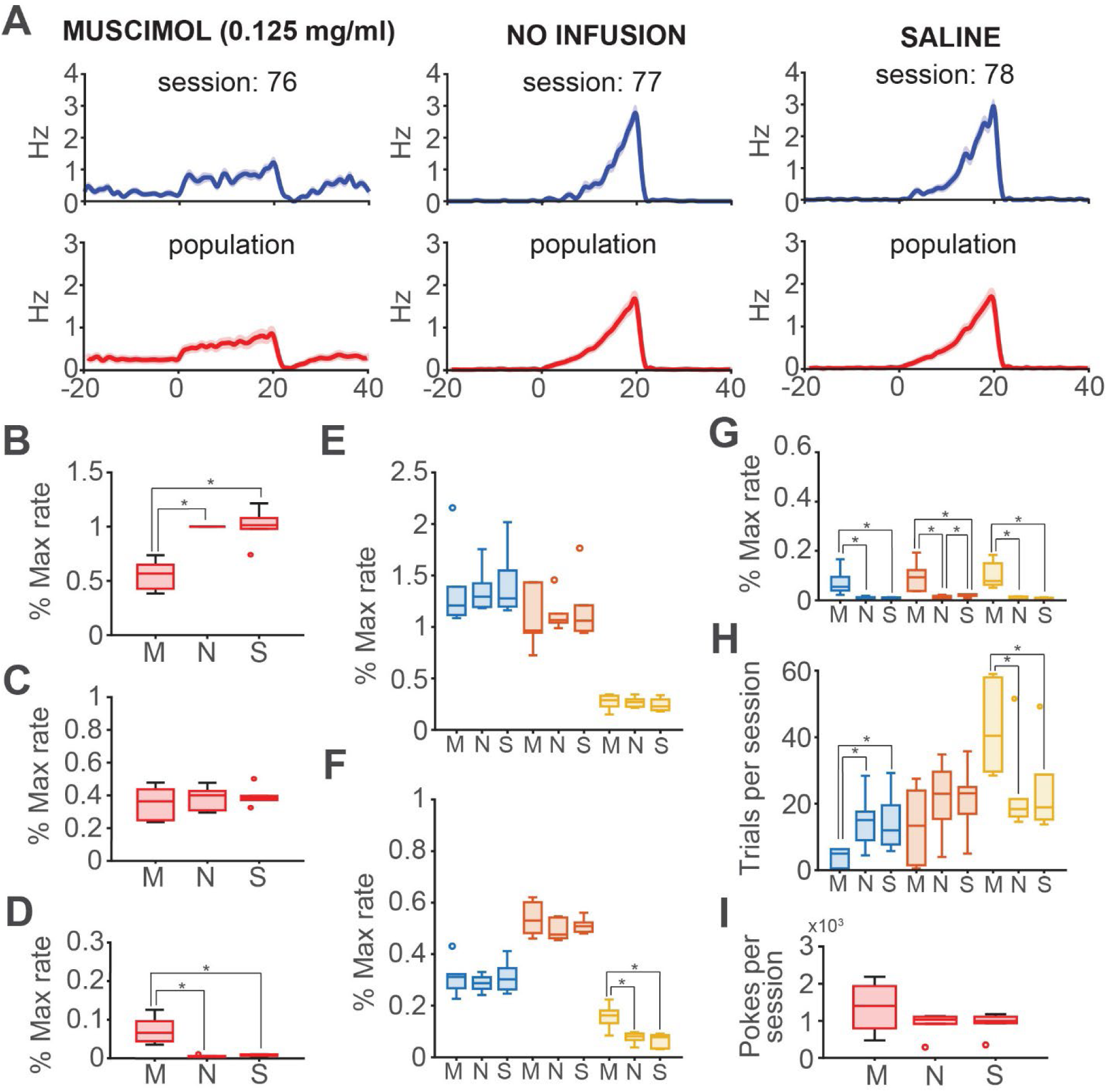
Muscimol-induced VMT inactivation reduces precision in 20s IT. **A.** (Top) Example average response curves for muscimol, no infusion, and saline sessions. (Bottom) Population average response curves. **B.** Muscimol decreases peak response rate. **C.** There are no changes in mean cued response rates. **D.** Muscimol increases mean pre-cue response rate. **E.** There are no significant differences in peak response rates per cluster. **F.** Muscimol increases mean response rate only in the RTR cluster trials. **G.** Muscimol increases mean pre-cue response rates in all the clusters. **H.** Muscimol decreases the number of OTR trials per session and increases the number of RTR trials per session. **I.** There are no differences in the average number of pokes per session between experimental groups.

Using the centroids corresponding to the k-means of the training trials, we clustered all experimental trials from the 20s IT muscimol, saline, and no infusion sessions. Contrasting with the significant decrease in average peak response rate, VMT inactivation had no significant effects when looking at peak response rates within behavioral clusters (OTR – muscimol: µ = 1.36, no infusion: µ = 1.38, saline: µ = 1.41; ETR – muscimol: µ = 1.44, no infusion: µ = 1.12, saline: µ = 1.17; RTR – muscimol: µ = 0.27, no infusion: µ = 0.27, saline: µ = 0.24) (Figure 4E). Mean cued response was increased in RTR muscimol trials (muscimol: µ = 0.16, no infusion: µ = 0.076, saline: µ = 0.066; Wilcoxon sign-ranked: p = 0.031) (Figure 4F), and mean pre-cue response rates were significantly increased in OTR (muscimol: µ = 0.072, no infusion: µ = 0.0062, saline: µ = 0.066; Wilcoxon sign-ranked: p = 0.0069), ETR (muscimol: µ = 0.096, no infusion: µ = 0.012, saline: µ = 0.0189; Wilcoxon sign-ranked: p = 0.031), and RTR muscimol trials (muscimol: µ = 0.1, no infusion: µ = 0.0077, saline: µ = 0.0054; Wilcoxon sign-ranked: p = 0.031) (Figure 4G). No difference was observed in total pokes per session between experimental sessions (muscimol: µ = 1357, no infusion: µ = 912, saline: µ = 928) (Figure 4I), thus VMT inactivation was affecting the precision of the responses. Changes in response rates were not simply sedation or decreased behavioral output. Lastly, muscimol significantly reduced the number of OTR trials per session (muscimol: µ = 3.94, no infusion: µ = 14.94, saline: µ = 14.38; Wilcoxon sign-ranked: p = 0.031), and significantly increased RTR trials (muscimol: µ = 13.38, no infusion: µ = 21.64, saline: µ = 21.5; Wilcoxon sign-ranked: p = 0.031) (Figure 4H), which became the majority of trials per session. These results show that first: a combination of inactivation-induced changes specific to individual cluster trials can result in an incorrectly measured effect when looking at the non-clustered behavioral data; and second: the VMT is necessary for precise performance in IT.

### 80s interval timing

After completing all muscimol infusion and control sessions at 20s IT, rats started training at 80s IT (Figure 5A). The first session displayed a broader response curve similar to that of the first 20s session. Pre-cue responses were virtually absent, and response rate curves showed very low values during the initial seconds after cue onset (Figure 5B). Over several weeks, response rates accumulated near the reward time.

**Figure 5.**
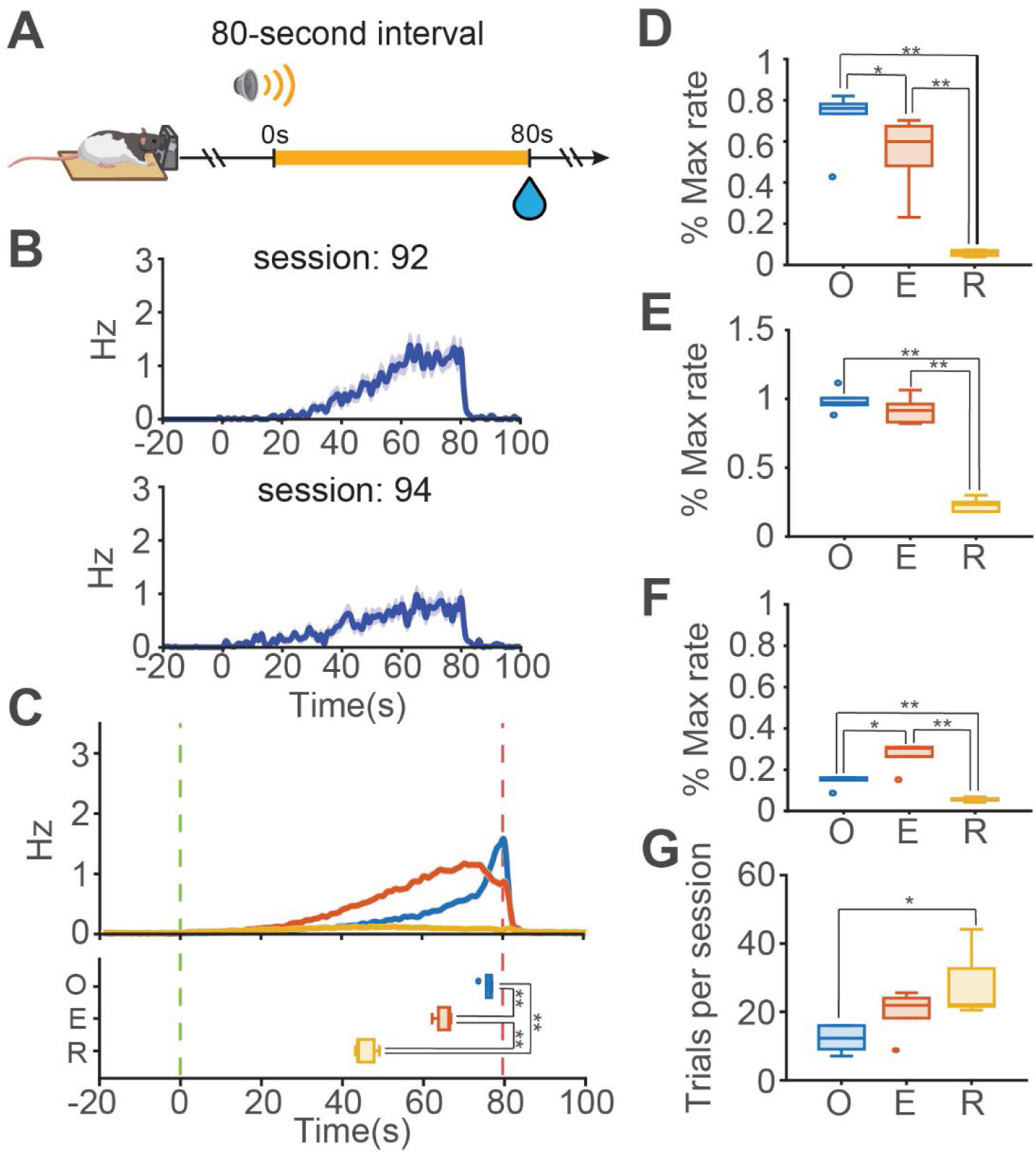
Timing at 80s interval. **A.** Schematic of the IT task using a 80s interval. **B.** Example session average response curves for 80s IT learning. Session 92 shows the first 80s IT session. **C.** (Top) Response clusters from all 80s IT trials. (Bottom) Time of peak response per cluster. On-Time Response (O) peaks at the time of expected reward; Early Timing Response (E) peaks before the expected reward; Random Timing Response (R) peaks earlier than both OTR and ETR. As in 5s and 20s IT each cluster is characterized by a pattern of response rate. *Cluster metrics:* **D.** Response rates at time of interval. **E.** Peak response rates. **F.** Mean cued response rates. **G.** Average number of trials per session.

We also clustered the 80s IT response trials, using a vector size of 120 (80s interval + 20s pre-cue + 20s post interval). The three characteristics clusters displayed at 5s and 20s were also present at 80s IT. OTR trials peaked near time of reward (µ = 75.9s, σ = 1.69s), ETR trials peaked sooner (µ = 66.03s, σ =3.28s), and RTR trials displayed a more variable peak response time (µ = 46.83s, σ = 2.5s) (Figure 5C). Response rates at the time of expected reward followed the same trend as at shorter intervals, also were significantly different between all three clusters (OTR: µ = 0.71, ETR: µ = 0.55, RTR: µ = 0.058; Wilcoxon ranksum: p = 0.0022) (Figure 5D). Average peak response rate was significantly higher in OTR and ETR trials compared to RTR trials (OTR: µ = 0.98, ETR: µ = 0.92, RTR: µ = 0.23; Wilcoxon ranksum: p = 0.0022) (Figure 5E). ETR trials showed a mean cued response rate significantly higher than OTR (OTR: µ = 0.14, ETR: µ = 0.27; Wilcoxon ranksum: p = 0.026) and RTR trials (RTR: µ = 0.056; Wilcoxon ranksum: p = 0.0022). Mean cued response in RTR trials was also significantly lower than OTR trials (Wilcoxon ranksum: p = 0.0022) (Figure 5F). Lastly, the number of trials in the RTR cluster was significantly higher than in OTR (OTR: µ = 13 trials, ETR: µ = 21.7 trials, RTR: µ = 18.9 trials; Wilcoxon ranksum: p = 0.0043) and not different from ETR (Figure 5G). This suggests that at longer intervals (80s IT single sessions last >2hrs) rats could be losing engagement with the task, thus having more trials with random responses.

The relative characteristics of each cluster remained stable across 5s, 20s, and 80s, supporting the existence of discrete behavioral strategies per trial. OTR trials show on average better timing with a delay before increasing responses; ETR trials show less behavioral inhibition, with a ramp up occurring sooner in the trial, peaking, and slightly lower at the time of expected reward; RTR trials most likely include trials in which the rat lost engagement, became distracted, failed trials, and in general low-responding trials.

### VMT inactivation in 80s IT

After the 80s interval timing training we also conducted pre-session infusion of muscimol (0.125 mg/mL) targeting the VMT. As seen at 20s, VMT inactivation at 80s IT caused an increase in pre-cue response rates (muscimol: µ = 0.13, no infusion: µ = 0.015, saline: µ = 0.011) and a decrease in peak response rates (muscimol: µ = 0.55, no infusion: µ = 1.0, saline: µ = 0.95), all statistically significant (Wilcoxon sign-ranked: p = 0.031) (Figure 6A-D) suggesting a decrease in the precision of the temporal response at this longer interval.

**Figure 6.**
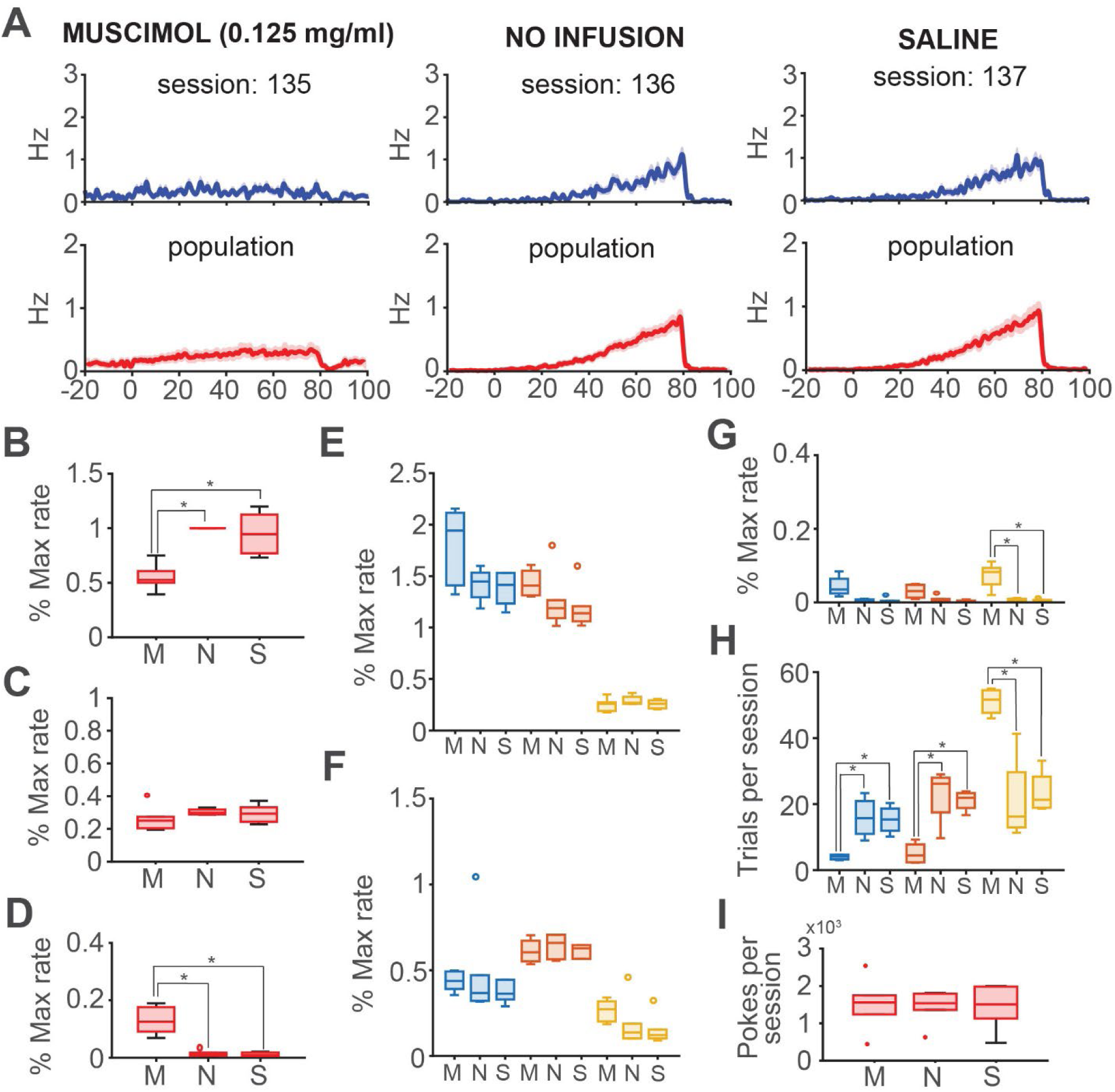
Muscimol-induced VMT inactivation reduces precision in 80s IT. **A.** (Top) Example average response curves for muscimol, no infusion, and saline sessions. (Bottom) Population average response curves. **B.** Muscimol decreases peak response rate. **C.** There are no changes in mean cued response rates. **D.** Muscimol increases mean pre-cue response rate. **E.** There are no significant differences in peak response rates per cluster. **F.** There are no differences in mean response rate. **G.** Muscimol increases mean pre-cue response rates in RTR trials. **H.** Muscimol decreases the number of OTR and ETR trials per session and increases the number of RTR trials per session. **I.** There are no differences in the average number of pokes per session between experimental groups.

We also clustered all experimental trials at 80s IT muscimol, saline, and no infusion sessions. Once again, VMT inactivation had no significant effects when looking at peak response rates within behavioral clusters (OTR – muscimol: µ = 1.79, no infusion: µ = 1.42, saline: µ = 1.69; ETR – muscimol: µ = 1.43, no infusion: µ = 1.26, saline: µ = 1.19; RTR – muscimol: µ = 0.25, no infusion: µ = 0.29, saline: µ = 0.26) (Figure 6E). Mean cued response rates were not different among conditions in each cluster (OTR – muscimol: µ = 0.24, no infusion: µ = 0.2, saline: µ = 0.21; ETR – muscimol: µ = 0.34, no infusion: µ = 0.35, saline: µ = 0.34; RTR – muscimol: µ = 0.13, no infusion: µ = 0.075, saline: µ = 0.064) (Figure 6F), and mean pre-cue response rates were significantly increased only in RTR muscimol trials (muscimol: µ = 0.073, no infusion: µ = 0.0074, saline: µ = 0.0054; Wilcoxon sign-ranked: p = 0.031) (Figure 6G). No difference was also observed in 80s IT in total pokes per session between experimental sessions (muscimol: µ = 1514, no infusion: µ = 1443, saline: µ = 1431) (Figure 6I), showing that VMT inactivation was affecting the precision of the responses. Lastly, the numbers of OTR and ETR trials per session were decreased in the muscimol sessions (OTR – muscimol: µ = 1.58, no infusion: µ = 8.06, saline: µ = 7.54; ETR – muscimol: µ = 2.62, no infusion: µ = 14.2, saline: µ = 15.4; Wilcoxon sign-ranked: p = 0.031), and significantly increased in RTR muscimol trials (muscimol: µ = 55.8, no infusion: µ = 37.8, saline: µ = 37.1; Wilcoxon sign-ranked: p = 0.031) (Figure 6H). The consistency of the results in both time intervals adds more evidence suggesting that rats performing IT may display multiple behavioral strategies. Strategies that are characterized by the precision of time estimation. This precision is impaired by VMT inactivation suggesting this structure as necessary for precise performance in IT.

## DISCUSSION

In this study we explored the role of the VMT in interval timing, more specifically in the retrieval of temporal memories. We found that inactivating the VMT using muscimol infusions results in a significant decrease in the precision of the retrieved memory. In the context of our behavioral paradigm, precise retrieval implies that most instrumental responses occurred during the cued interval with a high peak response. This response pattern, as seen in the normal training at 5s, 20s, and 80s, results in very low pre-cue response rates, that significantly increase under VMT inactivation. This loss of precision resembles the behavioral effects of mPFC inactivation in an interval timing task conducted by Buhusi et al. (2018). They report a significant muscimol-induced increase in responding during the ITI, which resulted in a lower elevation ratio of trial vs ITI average. They also report the flattening of the characteristic IT curve, an effect also reported after mPFC lesioning (Dietrich & Allen, 1998; Dietrich et al., 1997). We observed a clear flattening of the response curve denoted by a decrease of the average peak response rate (we discuss later how our results show this could result from an averaging artifact). mPFC inactivation has been also shown to impair interval discrimination even with large long/short interval ratios (Kim et al., 2009). Testing the role of the VMT in a time discrimination task could aid in comparing thalamic and mPFC involvement in this temporal ability and shed light on how timing information is processed and transmitted between these two areas.

The effects of our VMT inactivation were consistent throughout sessions and across subjects. Nevertheless, infusions lack the specificity of other techniques in establishing accurate targeting, or drug reach, as well as introducing the variability of individual-specific response to the infused drug. In this study we used a muscimol concentration of 0.125 mg/mL that in some rats produced a total loss of engagement with the task. For these animals we lowered the muscimol concentration to 0.0625 mg/m, and the rats were able to perform the task.

Histological analysis in our rats revealed that muscimol could have also impacted the submedius nucleus of the thalamus (SMT). This brain area has been associated with ascending nociception signaling from the spinal cord and has been shown important in the depression of persistent inflammatory pain (Tang et al., 2009; Wang et al., 2005; Xiao et al., 2005), it is necessary for updating Pavlovian conditioning (Alcaraz et al., 2015), and it participates in maternal behavior (Tasaka et al., 2026). The SMT has not been associated with timing or memory retrieval, and the effects of VMT inactivation were consistent across all animals. Nevertheless, a projection specific inactivation approach is necessary to further isolate VMT specific contributions to IT and avoid any unspecific inactivation.

The results of our clustering analysis suggest the presence of multiple behavioral strategies at play when the rat faces a single interval timing trial. The RTR cluster most likely encompasses all failed trials (if any), or otherwise those with a lack of the rat engaging with the task, being distracted, etc. These trials showed, across all trained intervals, the lowest mean and peak response rates. OTR trials point to a more accurate retrieved temporal memory, in which instrumental responses are delayed in time for a future reward. On the other hand, ETR trials display an early ramp up in instrumental responses, which peaks and begins trending down earlier than the rewarded interval. This decrease in response rate suggests decreasing behavioral effort due to a lack of accurate timing, or an underestimation in elapsed time. Thus, OTR and ETR trials, even with differences in accuracy, still show the animal is measuring time.

It is important to consider that our task cannot test the “real” retrieved time, which could be measured performing a peak interval (PI) timing task (Roberts, 1981). In a PI paradigm, the absence of reward in some trials prevents the rat from stopping their instrumental responses to consume the reward, thus the timing of the average peal response can be interpreted as the “real” retrieved time (Gibbon & Church, 1990; Meck, 2006; Meck et al., 1984). The response clusters we here report could be further investigated using this task, which could clarify how the rat’s effort is organized per trial in the absence of reward.

The existence of the response clusters also helps contextualize muscimol-induced changes in IT. VMT inactivation, besides increasing pre-cue response rates, resulted in a significant decrease in peak response. The clustered data reveals that these changes in averaged response rate are not the result of less precise instrumental responses. Peak response rate in OTR and ETR trials are not different in muscimol vs controls sessions. The underlying change is the number of trials per session in each cluster. The significant increase in the number of RTR trials and a decrease in the number of OTR trials (and ETR trials at 80s), results in the observed downward change in averaged peak response rate. Clustering reveals that the change in precision arises from loss of time-precise trials, and not a decrease in precise instrumental responding overall. OTR and ETR trials still display higher peak and mean cued response rates than RTR trials; thus, the behavioral strategies observed during training can still be displayed under VMT inactivation.

In the brain, these putative behavioral strategies most likely represent distinct underlying patterns of neural activity and communication between regions. Although there are no previous reports using a similar trial clustering approach in interval timing, there is sufficient behavioral and physiological data available to allow for individual groups to re-analyze and test our observations. Overall, our results show that the VMT is critical for retrieving precise temporal memories. This role in temporal memory processing is consistent with that of other areas in the mPFC-VMT-HC circuit of memory.

## Notes

### Competing Interest Statement

The authors have declared no competing interest.

### Funding

- National Institute of Environmental Health Sciences: T32ES033955

